# Characterization of the phototoxicity, chemigenetic profile, and mutational signatures of the chemotherapeutic CX-5461 in *Caenorhabditis elegans*

**DOI:** 10.1101/2019.12.20.884981

**Authors:** Frank B. Ye, Akil Hamza, Tejomayee Singh, Stephane Flibotte, Philip Hieter, Nigel J. O’Neil

**Affiliations:** Michael Smith Laboratories, University of British Columbia, Vancouver, V6T 1Z4, Canada; Department of Zoology, University of British Columbia, Vancouver, V6T 1Z4, Canada

## Abstract

New anti-cancer therapeutics require extensive *in vivo* characterization to identify endogenous and exogenous factors affecting efficacy, to measure toxicity and mutagenicity, and to determine genotypes resulting in therapeutic sensitivity or resistance. We used *Caenorhabditis elegans* as a platform with which to characterize properties of anti-cancer therapeutic agents *in vivo*. We generated a map of chemigenetic interactions between DNA damage response mutants and common DNA damaging agents. We used this map to investigate the properties of the new anti-cancer therapeutic CX-5461. We phenocopied the photoreactivity observed in CX-5461 clinical trials and found that CX-5461 generates reactive oxygen species when exposed to UVA radiation. We demonstrated that CX-5461 is a mutator, resulting in both large copy number variations and a high frequency of single nucleotide variations (SNVs). CX-5461-induced SNVs exhibited a distinct mutational signature. Consistent with the wide range of CX-5461-induced mutation types, we found that multiple repair pathways were needed for CX-5461 tolerance. Together, the data from *C. elegans* demonstrate that CX-5461 is a multimodal DNA damaging agent with strong similarity to ellipticines, a class of antineoplastic agents, and to anthracycline-based chemotherapeutics.

## INTRODUCTION

Cancer is driven by genetic and environmental factors. These factors can enhance or suppress the efficacy of anti-cancer therapeutics. Key to the adoption of targeted anticancer therapies is an understanding of the interactions between therapeutic agents, the tumour genotype and the tumour environment. For example, many anti-cancer therapeutics are mutagenic either as their primary mode-of-action or as a side-effect and some undergo xenobiotic metabolism to form new metabolites with different properties from the original drug [1,2]. For these reasons, the properties of anti-cancer therapeutics need to be assayed to determine: 1) mechanisms of action, 2) genetic alterations affecting therapeutic sensitivity and resistance, 3) mutagenicity and 4) the effect of environment on the therapeutic agent.

Chemotherapeutic genotoxicity and efficacy can be dependent on genotype. Screening for therapeutic-sensitive and -resistant genotypes can identify tumour-specific genetic biomarkers and add to the understanding of the mechanism of anti-cancer therapeutics. Many pharmacogenomic screens have been conducted in human cell lines using RNAi, or more recently, CRISPR sgRNA libraries, to identify anti-cancer chemi-genetic interactions [3–7]. While useful, the genotypes of many cell lines are not well characterized and can evolve because of genomic instability. A more genetically stable, near isogenic assay system could be utilized to take a more reductive approach to determine chemi-genetic interactions with new potential therapeutics.

The nematode *Caenorhabditis elegans* is an attractive animal model with which to characterize the properties of anticancer therapeutics. The small size, ease of handling, and powerful genetic tools of *C. elegans* provide a sophisticated *in vivo* platform that combines the technical advantages of a microorganism with greater biological complexity. *C. elegans* has been used to screen for and characterize compounds affecting meiosis [8,9] and development [10]. *C. elegans* has also proven useful for determining mutational frequencies and signatures of DNA damage response (DDR) mutants and genotoxic agents [11–15].

We used *C. elegans* to characterize the *in vivo* properties of CX-5461, a promising anti-cancer therapeutic currently in clinical trials [16,17]. CX-5461 was first described as an orally bioavailable RNA Pol I inhibitor that exhibited anti-tumour activity in murine xenograft models [18] and was the first RNA Pol I inhibitor to be tested in clinical trials [17]. CX-5461 also stimulates ATM/ATR activation [19], and rapamycin-associated signaling [20]. More recently, it was found that homologous recombination deficient cancer cells are sensitive to CX-5461 and that this sensitivity may be due to the stabilization of G-quadruplex forming DNA structures that could affect DNA replication [21]. This has led to a clinical trial focusing on patients with homology-directed repair (HDR)-deficient tumours [16]. However, the mechanisms underlying the tumour cell killing and the *in vivo* properties of CX-5461 are still unclear.

We assayed CX-5461-mediated photosensitivity, mutagenicity, mutational signatures, and identified genotypes that are sensitive to CX-5461. We found that CX-5461 is a multimodal genotoxic agent with similarities to antineoplastic ellipticine and anthracycline-based anti-cancer agents. A better understanding of CX-5461 can lead to more informed treatment approaches.

## RESULTS

### CX-5461 is a photosensitizer that generates ROS upon exposure to UVA

Photosensitivity is a common side-effect of many therapeutics [22]. Clinical trials evaluating CX-5461 in patients with hematologic or advanced solid tumors reported cases of photosensitivity [16,17]. We used *C. elegans* as an *in vivo* model to investigate the photosensitivity of CX-5461. We focused on the effect of UVA radiation on CX-5461 for several reasons: 1) CX-5461 absorbs UVA and UVB radiation, 2) other quinolone-based molecules can trigger photosensitivity upon UVA irradiation [22], 3) UVA passes through clouds and glass, accounting for more than 90% of the UV radiation reaching the Earth’s surface, and 4) UVA penetrates deep into the dermis and triggers chemical-induced photosensitivity.

First, we attempted to re-create the CX-5461-induced photosensitivity in wild-type *C. elegans.* Young adult animals were exposed to CX-5461 for ~16 hours and then exposed to UVA radiation. Photosensitivity was measured by assessing the viability of F1 progeny from exposed animals. Wild-type animals were not sensitive to CX-5461 or UVA alone but were sensitive to CX-5461 + UVA exposure (Figure 1A). Increasing either the concentration of CX-5461 or the amount of UVA radiation enhanced the cytotoxicity in a dose dependent manner (Figure 1A). To assess if the photosensitivity was limited to the germline, we assayed CX-5461 photosensitivity in L1 larvae. Synchronized L1 larvae were arrested by starvation and treated with 100 μM CX-5461 for ~16 hours followed by exposure to 300 J/m^2^ UVA radiation. Photosensitivity was measured by assessing the developmental stage of the population after 96 hours. L1 wild-type animals were not sensitive to CX-5461 or UVA alone but were sensitive to CX-5461 + UVA exposure with many animals failing to develop to the adult stage (Figure 1B). To test if CX-5461 photosensitivity was conserved in other species, we assayed UVA-mediated CX-5461 photosensitivity in mismatch repair defective and proficient human cancer cell lines (HCT116 and HT29, respectively) and in wild-type and homologous recombination defective budding yeast (*Saccharomyces cerevisiae*). Both human colorectal cancer cell lines exhibited UVA-induced dose-dependent CX-5461-mediated photosensitivity (Figure 1C). Similarly, wild-type and *rad52* yeast also exhibited dose-dependent CX-5461-mediated photosensitivity (Figure 1D).

**Figure 1.**
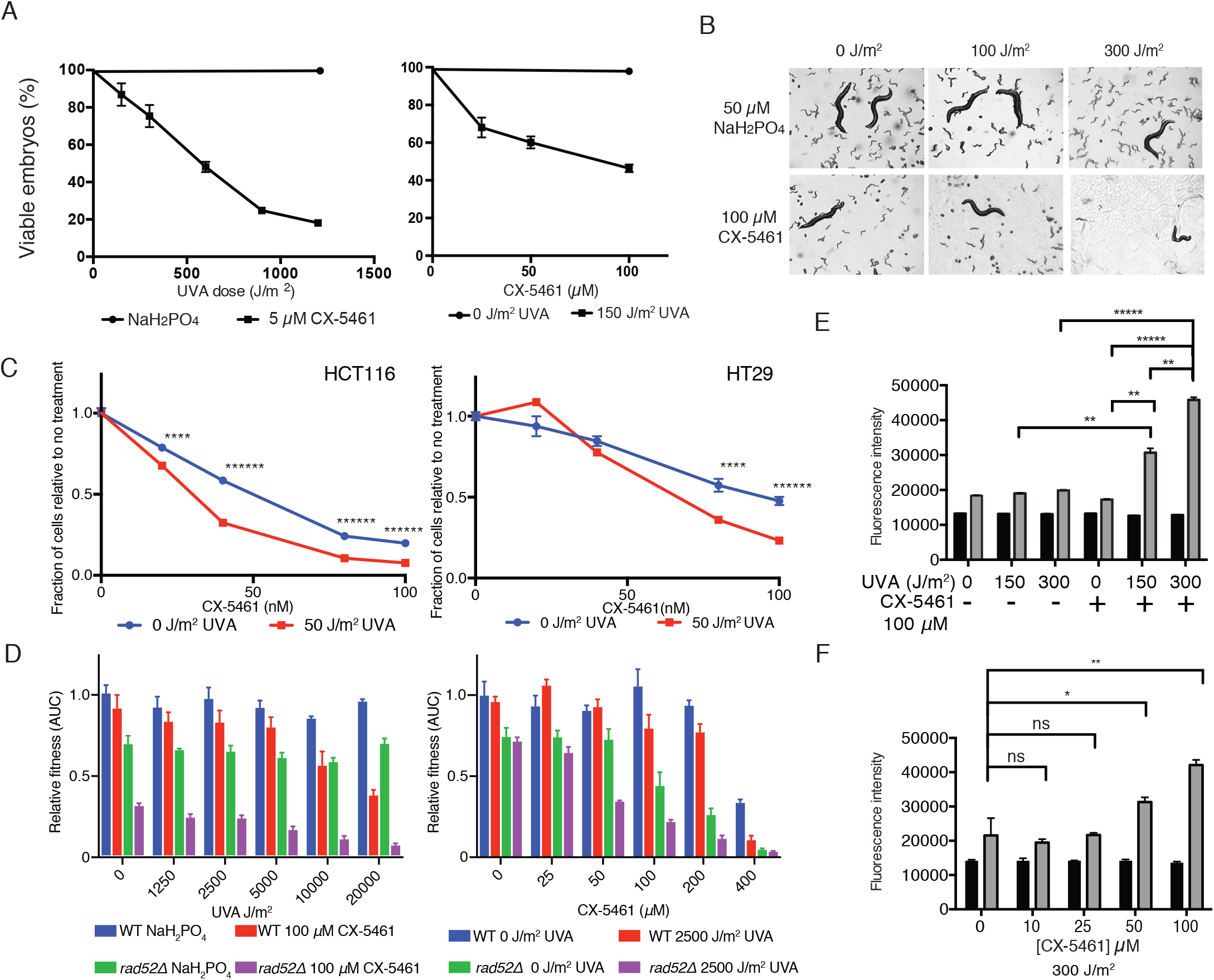
CX-5461 is a photosensitizer in *C. elegans,* human cancer cell lines and yeast. A. Viability of WT *C. elegans* embryos from adult animals exposed to CX-5461 and irradiated with UVA. Left-constant [CX-5461]; Right-constant UVA dose. B. Representative images of WT *C. elegans* populations 96 hours after CX-5461 +UVA exposure of synchronized WT L1 larvae. The large animals are the treated P0 individuals. C. HCT116 and HT29 colorectal cancer cell lines were treated with increasing concentrations of CX-5461 and exposed to UVA irradiation in 96-well format and cell nuclei counted after 96 hours. Student’s t-test ****, P< 0.0005; ******, P<0.000005. D. Growth curve analysis of the relative fitness of wild-type and rad52 yeast exposed to CX-5461 + UVA radiation. Left-fixed [CX-5461]; Right-fixed UVA dose. E. Intracellular ROS levels were measured in CX-5461 + UVA treated wild-type *C. elegans* with constant [CX-5461]. F. Intracellular ROS levels were measured in CX-5461 + UVA treated wild-type *C. elegans* with a constant UVA dose. Student’s t-test *, P< 0.05; **, P<0.005, *****, P<0.000005.

The phototoxicity of some fluoroquinolones can be attributed to the generation of reactive oxygen species (ROS) after exposure to UVA radiation [23]. To determine whether CX-5461 generated ROS upon UVA radiation, we used 2’,7’-dichlorodihydrofluorescein diacetate (H_2_DCFDA) as an intracellular fluorescent probe to measure ROS [24] in CX-5461 + UVA exposed *C. elegans.* We observed a significant dose-dependent ROS increase in worms treated with CX-5461 followed by UVA exposure (Figure 1E and 1F). Increasing UVA or CX-5461 increased the amount of ROS produced (Figure 1E and 1F). Taken together, these data suggest that CX-5461 is a photosensitizer that results in cytotoxicity due to the production of ROS.

### CX-5461 exposure results in SNVs and GCRs

CX-5461 has been shown to stabilize G-quadruplexes [21]. G-quadruplex stabilization can cause replication-associated mutagenic events [11,25–27]. To characterize the frequency and spectrum of mutagenic events induced by CX-5461 and CX-5461 + UVA, we used a genetic balancer in wild-type *C. elegans* to capture and characterize CX-5461-induced lethal mutations in the presence and absence of UVA. The *eT1* balancer is a reciprocal translocation of approximately half of chromosome III and half of chromosome V and can capture both single nucleotide variants (SNVs) and copy number variants (CNVs) in balanced regions, including terminal deletion events and translocations [15].

Exposure to CX-5461 or CX-5461 + UVA resulted in high frequencies of balanced lethal mutations and dominant sterile F1 animals, which produced no progeny (Figure 2A). Four balanced recessive lethal mutations were recovered from a screen of 200 F1 progeny from individuals treated with 100 μM CX-5461. UVA radiation increased the mutagenicity of CX-5461 more than four-fold. Nineteen balanced recessive lethal mutations were recovered from a screen of 200 F1 progeny from individuals treated with 100 μM CX-5461 + 100 J/m^2^ of UVA radiation.

**Figure 2.**
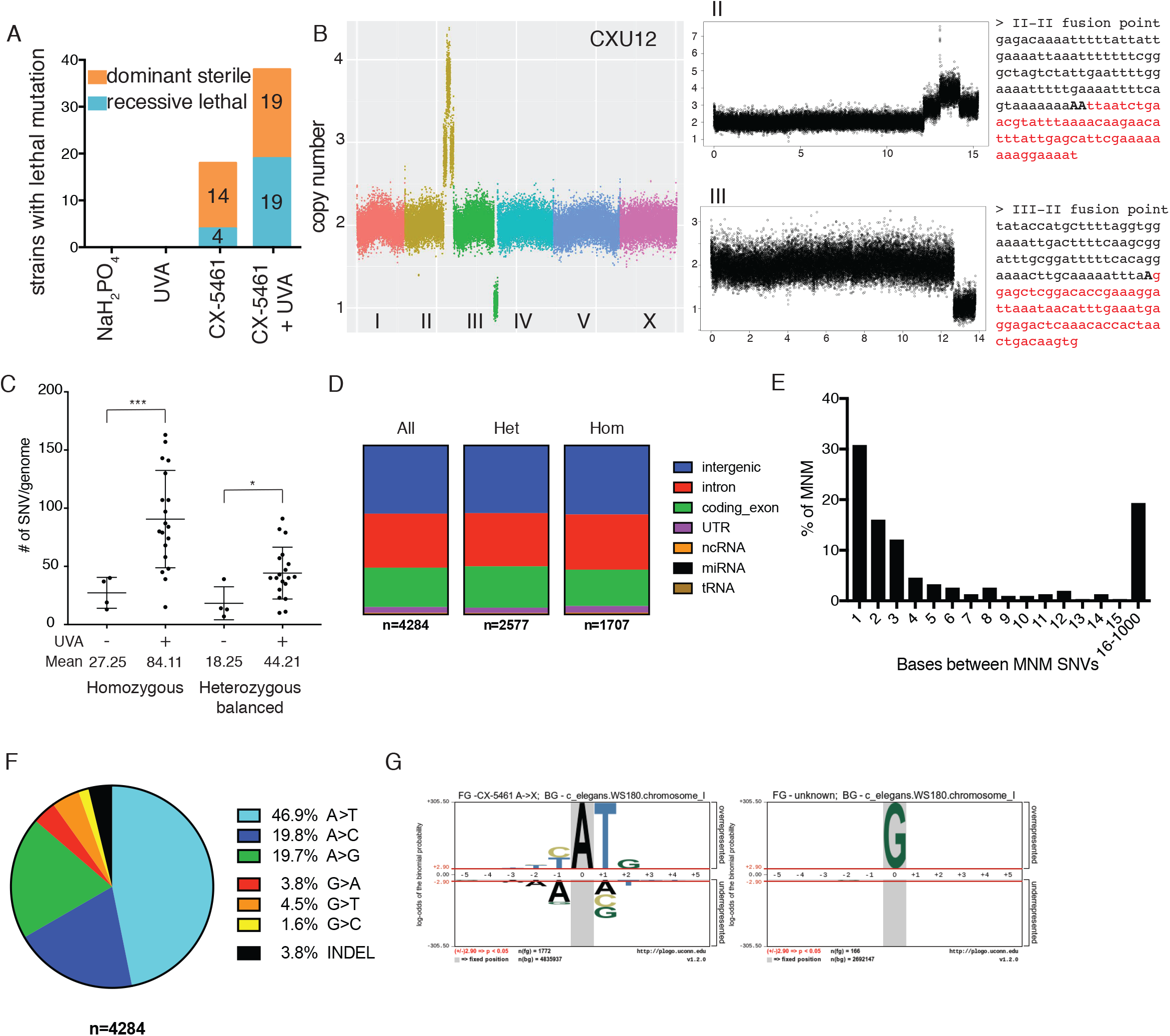
Exposure to CX-5461 or CX-5461 + 100 J/m2 UVA results in high frequencies of mutations. A. Number of balanced recessive lethal mutations and dominant sterile mutations. n=200 for each condition. B. Coverage plot of CX-5461 + UVA-induced genome rearrangements in sample CXU12. Whole genome (Left). Detailed coverage plot of chromosome II (Top right) and chromosome III (bottom right). Sequence at the fusion shown on right. Microhomology in bold. C. Number of homozygous and heterozygous balanced SNVs/genome. Welch’s t-test *** p<0.0005, * p<0.05 D. Distribution of SNVs in coding and non-coding elements. E. Distance between SNVs in multi-nucleotide mutations. F. SNV mutational signature of CX-5461. G. pLOGO of extended sequence context of CX-5461-induced SNVs.

To elucidate the mutational signatures of CX-5461 and CX-5461 + UVA, we sequenced the genomes of the 23 strains with *eT1-*balanced lethal mutations. The CX-5461-treated genomes contained a range of mutation types, including large copy number variations (CNVs) and single nucleotide variations (SNVs). First, we analyzed the mutations in the balanced regions to identify the lesions responsible for the lethal phenotype. In the CX-5461 mutated strains, 9/23 contained large copy number variations in the balanced regions that could account for the lethality (Table 1; Supplemental Figures 1-2) and 14/23 strains contained SNVs in essential genes (Table 1).

**Table 1:** CX-5461-induced SNVs and CNVs

|  | Line | SNVs | Balanced heterozygous SNVs | Homozygous SNVs | Balanced CNVs | Putative lethal mutation |
| --- | --- | --- | --- | --- | --- | --- |
| CX-5461 | 1 | 68 | 14 | 52 | III Del |  |
|  | 2 | 58 | 5 | 51 | V Del |  |
|  | 3 | 348 | 47 | 11 |  | *chc-1* stop |
|  | 4 | 46 | 13 | 19 |  | *F54C8.4* stop |
| CX-5461 + UVA | 1 | 159 | 38 | 62 |  | *plrg-1* FS |
|  | 2 | 283 | 52 | 117 | III Del |  |
|  | 3 | 190 | 34 | 107 |  | *mrpl-1* |
|  | 4 | 241 | 60 | 130 | III Del | *strd-1/mlc-7* |
|  | 5 | 144 | 37 | 80 | V Del |  |
|  | 6 | 258 | 57 | 143 | III Dp | multiple |
|  | 7 | 178 | 45 | 87 |  | *hpo-26* |
|  | 8 | 121 | 35 | 68 | III Del/Inv |  |
|  | 9 | 179 | 52 | 95 | V Del III Inv | multiple |
|  | 10 | 201 | 33 | 107 |  | T05H4.10 |
|  | 11 | 151 | 35 | 74 | V Dp | *npp-16* |
|  | 12 | 138 | 23 | 56 | III Del |  |
|  | 13 | 54 | 11 | 15 | V trans |  |
|  | 14 | 485 | 89 | 157 | V Del | *let-413* |
|  | 15 | 243 | 43 | 84 |  | *klp-7* |
|  | 16 | 154 | 22 | 39 |  | *pri-1* |
|  | 17 | 222 | 39 | 79 |  | *ncx-2* |
|  | 18 | 168 | 29 | 43 | III Del |  |
|  | 19 | 195 | 10 | 31 |  | None |

Del-Deletion Dp-Duplication Inv-Inversion Trans-Translocation

### Analysis of CX-5461-induced CNVs

Most CX-5461- and CX-5461 + UVA-treated genomes contained at least one CNV CNVs ranged from simple deletions to complex events involving deletions, duplications, and translocations (Supplemental Figure 1 and 2). The high frequency of CX-5461-induced CNVs was consistent with the observation that DNA double strand break repair genes, such as the homology-directed repair (HDR) gene *brc-2* and the microhomology mediated end-joining (MMEJ) gene *polq-1,* are required for CX-5461 tolerance in *C. elegans* [21]. CNV breakpoints frequently contained regions of microhomology consistent with MMEJ (Table 1; Figure 2B). Analysis of the regions surrounding the CNV breakpoints found DNA repeats (simple, tandem, and inverted) flanking some of the breakpoints.

**Table 2:**
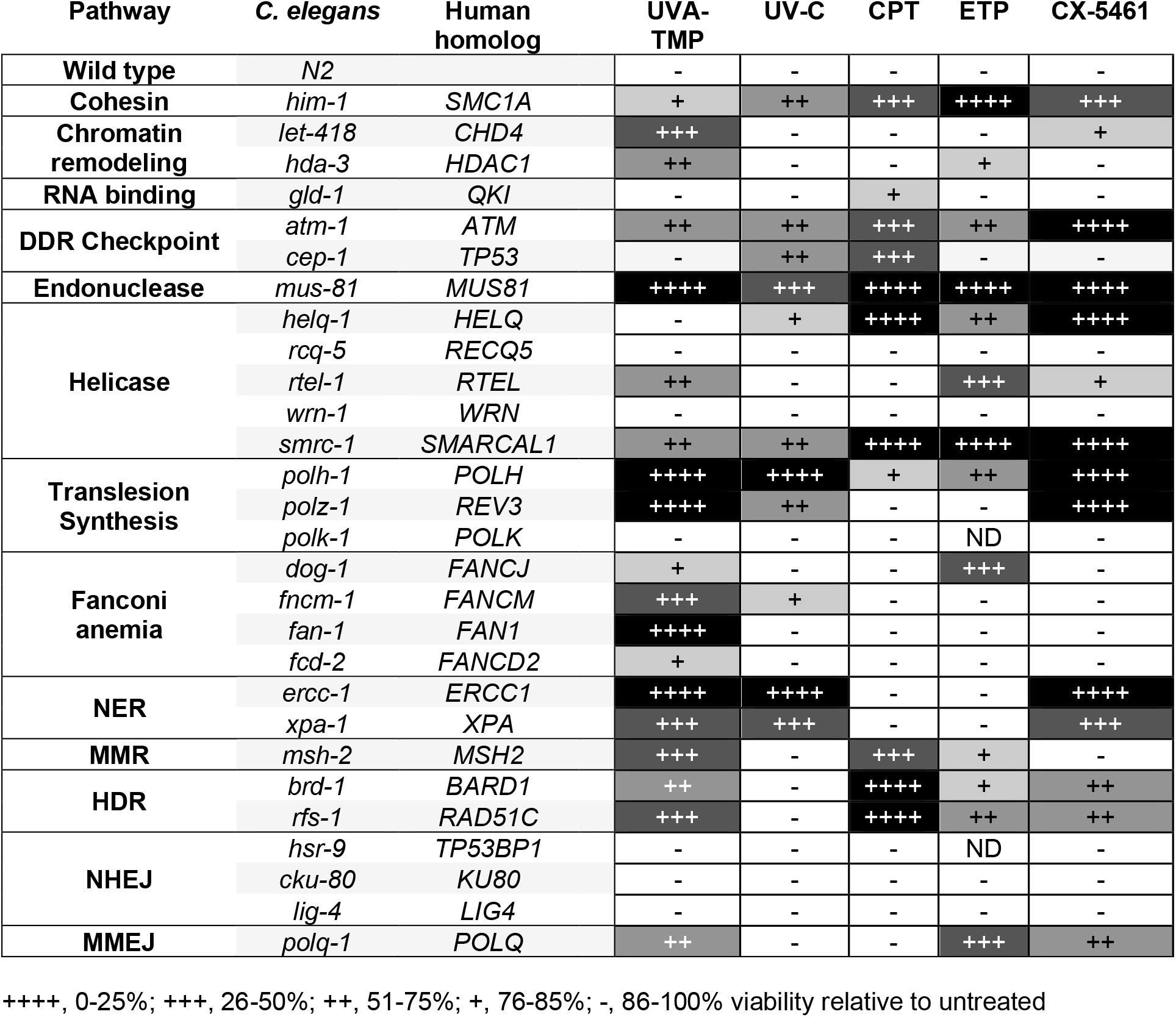
Chemigenetic profiles of *C. elegans* DNA damage response mutants.

### Analysis of CX-5461-induced SNVs

All CX-5461-exposed genomes contained a high frequency of heterozygous and homozygous SNVs (Table 1). Genomes exposed to CX-5461 + UVA had more homozygous and heterozygous SNVs in the balanced region compared to those exposed to CX-5461 alone (Figure 2C). The increased frequency of SNVs in the CX-5461 + UVA treated genomes was consistent with the increased frequency of balanced lethal mutations.

All 4,284 SNVs were included in the analysis because there were no obvious differences in the characteristics of the mutational profiles of heterozygous or homozygous SNVs or between the CX-5461 and CX-5461 + UVA-induced SNVs. The SNVs were distributed throughout the genome with no bias for coding or non-coding regions (Figure 2D) or chromosome location (Supplemental Figure 3; Supplemental Table 1). 517 SNVs (12%) were present in 212 multinucleotide mutation (MNM) clusters consisting of 2-13 SNVs within a 1000 bp region. More than 80% of the SNVs in MNMs were less than 15 bases from the neighboring mutations (Figure 2E). It was possible that the SNVs were the product of repair or bypass of CX-5461-stabilized G-quadruplexes, so we searched 100 bases 5’ and 3’ of each SNV for G-quadruplex forming structures using QuadBase2 webserver [28]. SNVs were not strongly correlated with G-quadruplex forming sequences. Only 0.75% of mutations flanking regions contained predicted G-quadruplexes compared to 0.45% in a control set of EMS mutations from the Million Mutation Project [29].

CX-5461-induced SNVs exhibited a distinct mutational signature. Greater than 80% of the SNVs were A to X changes with nearly 50% being A to T transversions (Figure 2F). To better understand the mutagenicity of CX-5461, we used the pLogo, a probability Logo generator, to examine the extended sequence context of the A to X mutations [30]. We observed changes in the frequency of bases both 5’ and 3’ of the mutated adenine. Most notably, 70% of the bases immediately 3’ (+1 position) of the mutated adenine were thymine. Guanine was overrepresented in the +2 position and cytosine was overrepresented in the −1 and −2 positions. In contrast, no extended sequence context was detected flanking mutated guanine (Figure 2G). Although there was a higher frequency of SNVs in the CX-5461 + UVA samples compared to CX-5461 genomes, we saw no difference in the types of SNVs suggesting that UVA exposure enhanced the frequency of CX-5461-induced SNVs but did not change the mutational mechanism.

To identify sequence motifs that may be more prone to CX-5461 mutagenesis, we looked for sites that were mutated in more than one line. Forty-seven sites were mutated in two or more lines (127 SNVs). We analyzed 100 bases flanking each of the frequently-mutated sites for sequences predicted to form secondary structures and found 25/47 flanking regions contained inverted or tandem repeats (53%). For comparison, a similar analysis of 3,719 regions flanking EMS-induced mutations from the Million Mutation Project [29] found 753 repeats (20.2%). From this, it appears that CX-5461-induced mutations are more common in regions containing tandem or inverted repeats.

### CX-5461 intercalates into DNA

The broad distribution of CX-5461-induced mutations suggested that CX-5461 can affect DNA even in the absence of G-quadruplex structures. Previous *in silico* analysis predicted that the pharmacophore of CX-5461 can intercalate into a crystal structure of DNA (PDB code 1Z3F) [31] in a manner similar to the antineoplastic agent ellipticine [32]. To test whether CX-5461 could intercalate into DNA, we incubated CX-5461 with a PCR-generated dsDNA and visualized the migration of DNA on a 1% agarose gel with the dsDNA-specific dye SYBR-Safe. Incubation of the dsDNA with CX-5461 resulted in a slower migrating DNA band suggesting that intercalation had occurred (Figure 3A). The disruption of dsDNA was greater when the DNA was denatured and reannealed in the presence of CX-5461. At higher concentrations, the DNA-CX-5461 complex did not migrate into the gel (Figure 3A).

**Figure 3.**
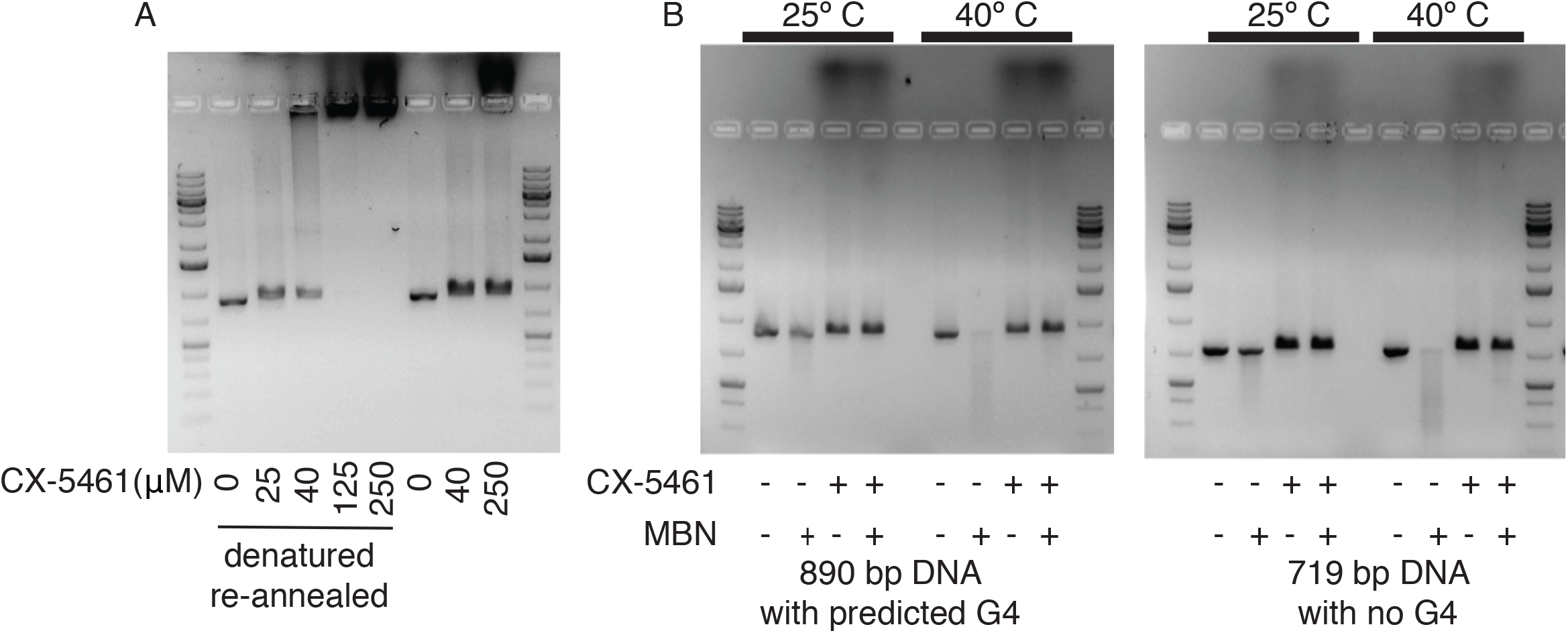
CX-5461 stabilizes DNA duplex structures. A. CX-5461 binds to and impedes the migration of dsDNA on a 1% agarose gel. CX-5461 binding is enhanced by DNA denaturation and re-annealing (Lanes 2-6). The effect was less in samples that were incubated without denaturation and reannealling (Lanes 7-9). B. CX-5461 stabilizes PCR products and the complex was more resistant to Mung Bean Nuclease (MBN) cleavage.

Intercalation of ellipticine into DNA results in partial unwinding and distortion of the DNA duplex [31]. To determine whether CX-5461 intercalation distorted DNA structure and whether G-quadruplex sequences were required, we incubated a PCR product predicted to form a G quadruplex (G4) and a PCR product without a predicted G quadruplex (non-G4) with mung bean nuclease (MBN), which cleaves single-stranded or distorted double-stranded DNA, for one hour at the specified temperature and assessed the endonuclease activity on a 1% Syber-Safe containing agarose gel. CX-5461 protected both G4 and non-G4 containing DNA fragments from MBN activity relative to DNA without CX-5461 (Figure 3B). At 40 □ both PCR products without CX-5461 were degraded, whereas the samples containing CX-5461 were not degraded suggesting that CX-5461 could increase the thermal stability of dsDNA.

### Genotypic sensitivities to CX-5461

The high frequency of both SNVs and CNVs suggested that the DNA damage response to CX-5461 is complex. To further characterize the DNA damage response to CX-5461, we used an acute exposure assay on a panel of 28 *C. elegans* DNA replication and repair mutants to generate a genetic sensitivity profile for CX-5461. We then compared the CX-5461 sensitivity profile to the profiles of the topoisomerase I poison camptothecin (CPT); the topoisomerase II poison etoposide (ETP); the interstrand crosslinking (ICL) agent UVA-activated trimethyl psoralen (UVA-TMP); and UV-C radiation (UV-C), which causes thymine dimers and photoproducts.

CX-5461 sensitivity was observed in 14 of the 28 DNA damage response mutants tested (Table 2). Mutations in genes implicated in replication stress *(mus-81, smrc-1, helq-1, brd-1, atm-1, polq-1, polz-1, polh-1, him-1)* resulted in sensitivity to CX-5461. The CX-5461 sensitivity profile was distinct from the other DNA damaging agents, sharing some but not all genotypic sensitivities. Of the 14 CX-5461 sensitive strains, 11 were sensitive to UVA-TMP, 9 were sensitive to UV-C, 8 were sensitive to CPT and 8 were sensitive to ETP. Given the overlapping sensitivities of CX-5461 and UVA-TMP, it was surprising that the Fanconi Anemia pathway mutants were not sensitive to CX-5461, demonstrating that CX-5461 does not result in ICLs. The CX-5461 sensitivity profile has similarity to that of the topoisomerase poisons CPT and ETP (Figure 4A). The main difference being that TLS and NER mutants were very sensitive to CX-5461 and not to either topoisomerase poison.

**Figure 4.**
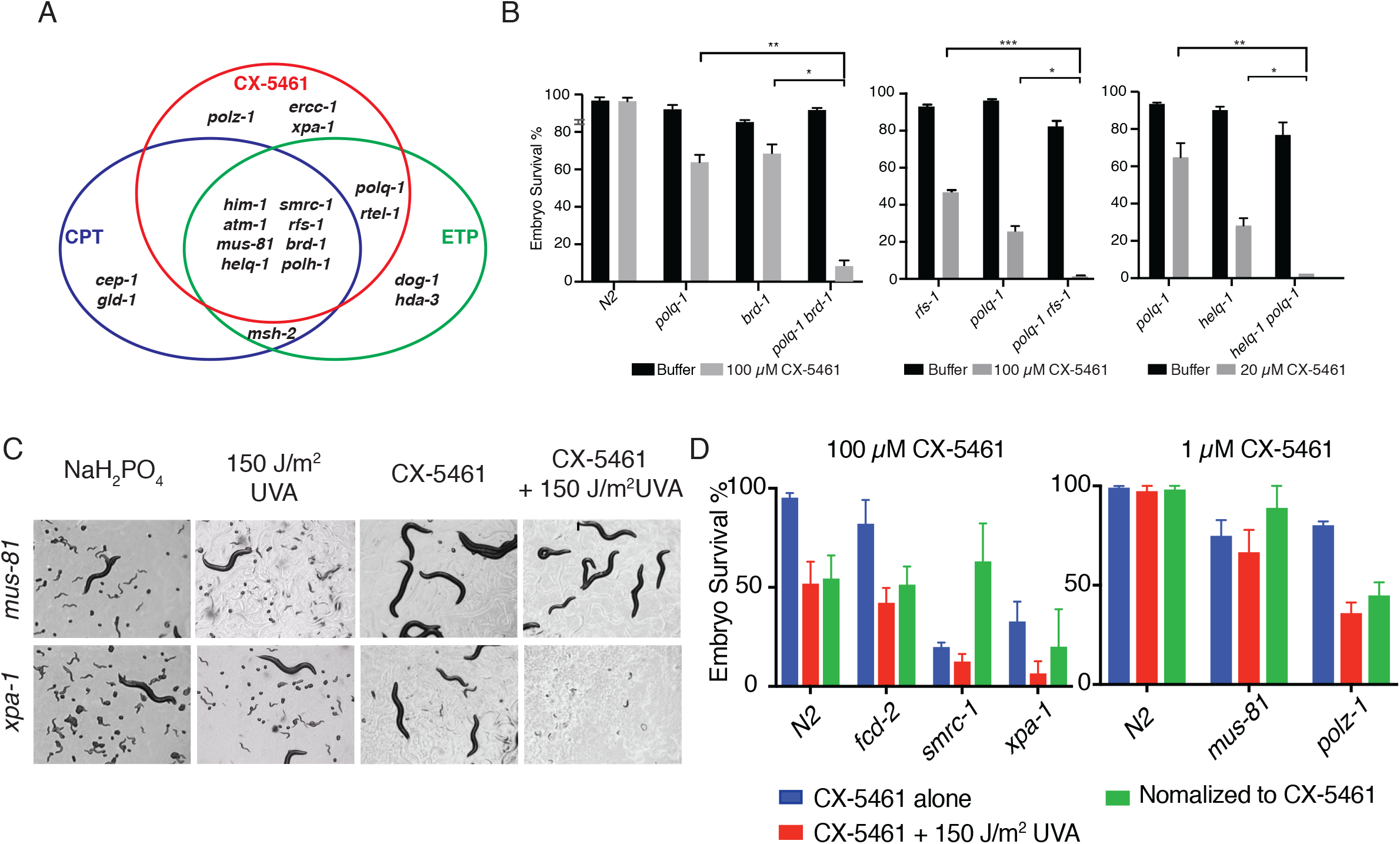
Genotypic sensitivity to CX-5461. A. Genotypic sensitivity profile of CX-5461. Venn diagram shows that the CX-5461-sensitive mutants also exhibited sensitivity to other DNA damaging agents, including the topoisomerase poisons camptothecin (CPT), and etoposide (ETP). B. Loss of *polq-1* sensitizes HDR-associated mutants (*brd-1, rfs-1*, and *helq-1*) to CX-5461. The bar graph showed the embryo survival rate for adult animals treated with the indicated dose of CX-5461. Student’s t-test *, P<0.05 **, P<0.005, *** P<0.0005. C. UVA enhances the toxicity of CX-5461. The image shows the growth and development of worms four days after L1 larva-treatment. Upon CX-5461 treatment and UVA irradiation, *mus-81* mutants developed into sterile adults, whereas *xpa-1* mutants arrested in L1. D. Differential sensitivity of worm mutants upon exposure to CX-5461 + UVA. CX-5461 hypersensitive mutants were tested at low [CX-5461] (Right). Note that *xpa-1* and *polz-1* are the only mutants that are more sensitive to CX-5461+UVA when normalized to account for the sensitivity to CX-5461 alone.

### HDR and MMEJ are required for CX-5461 tolerance

Mutations affecting the different double strand break repair pathways caused differential sensitivity to topoisomerase poisons and CX-5461. Non-homologous end joining (NHEJ) mutants were not sensitive to topoisomerase poisons or CX-5461. HDR mutants (*brd-1, rfs-1, helq-1*) were exquisitely sensitive to CPT but only mildly sensitive to ETP, whereas the MMEJ mutant *polq-1* was very sensitive to ETP but not sensitive to CPT. In contrast, mutations affecting either HDR or MMEJ resulted in moderate sensitivity to CX-5461. To test whether HDR and MMEJ were contributing independently to the repair of CX-5461 induced lesions, we tested *polq-1 brd-1, rfs-1 polq-1* and *helq-1 polq-1* double mutants for increased sensitivity to CX-5461. In all three cases, the double mutants exhibited increased CX-5461 sensitivity suggesting that HDR and MMEJ contributed independently to the repair of CX-5461-induced lesions (Figure 4B).

### Replication arrested nucleotide excision repair mutants are sensitive to CX-5461

Nucleotide excision repair mutants *xpa-1* and *ercc-1* were sensitive to CX-5461. NER repairs bulky single-stranded DNA lesions such as those formed by UV light and some cancer chemotherapeutics. Most NER activity is transcription-coupled (TC-NER). It is possible to assay the effect of DNA damaging agents on transcription-coupled repair by exploiting starvation-induced L1 diapause, in which replication is arrested. L1 larvae with TC-NER defects exposed to transcription-blocking DNA damaging agents are unable to reinitiate development when released from arrest [33], whereas, DNA damaging agents that do not block transcription reinitiate development. To test whether CX-5461 caused transcription-blocking lesions, we assayed CX5461 sensitivity in L1 arrested NER mutants. Replication-arrested L1 larvae were exposed to CX-5461 + UVA, released from arrest, and their development stages were assessed 96 hours later. CX-5461 + UVA-treated *xpa-1* and *ercc-1* L1 larvae failed to develop to later larval stages suggesting that CX-5461 can cause transcription-blocking lesions (Figure 4C). In contrast, the replication-associated CX-5461 hypersensitive mutant, *mus-81* could reinitiate development after L1 CX-5461 exposure and developed into sterile adults. These data demonstrate that CX-5461-induced lesions can block both transcription and replication.

### TLS and NER mutants exacerbate CX-5461 photosensitivity

To test whether the CX-5461-induced photosensitivity was due, in part, to increased DNA damage or changes in the nature of the DNA damage, we tested select mutants in the panel of *C. elegans* DNA replication and repair mutants for increased photosensitivity. Most DNA repair mutants were no more photosensitive to CX-5461 + UVA than wild-type animals (Figure 4D). However, the translesion polymerase mutant *polz-1* and the nucleotide excision repair mutant *xpa-1* exhibited greater embryonic death than expected. These results are consistent with the observation that CX-5461 generates ROS after UVA exposure (Figure 1E and 1F) and TLS and NER are required for the repair of DNA damage induced by ROS generated by UVA exposure [12].

### CX-5461 sensitive mutants and G-quadruplex stabilization

There are similarities between worms exposed to CX-5461 and worms lacking *C. elegans* FANCJ ortholog, *dog-1.* CX-5461 can stabilize G-quadruplexes [21] and the loss of DOG-1 results in the formation and/or stabilization of G-quadruplex structures [27]. Furthermore, CX-5461-exposed animals and *dog-1* mutants exhibit large and small chromosome rearrangements that often have MMEJ-signatures at the breakpoints [11,34]. To further investigate the similarities between CX-5461 exposure and loss of *dog-1,* we tested whether loss of *dog-1* resulted in negative genetic interactions with the CX-5461-sensitive mutants by measuring the viability of *dog-1* CX-5461-sensitive double mutants using a generational survival assay (Figure 5). The *polq-1* mutant was very sensitive to loss of DOG-1 with fewer than 50% of the lines surviving to the third generation. *mus-81* and *brd-1* mutants were also sensitive to *dog-1-*induced G-quadruplex stabilization. However, not all CX-5461 sensitive mutants exhibited genetic interaction with *dog-1* as the loss of *polz-1* did not affect the viability of *dog-1* mutants.

**Figure 5.**
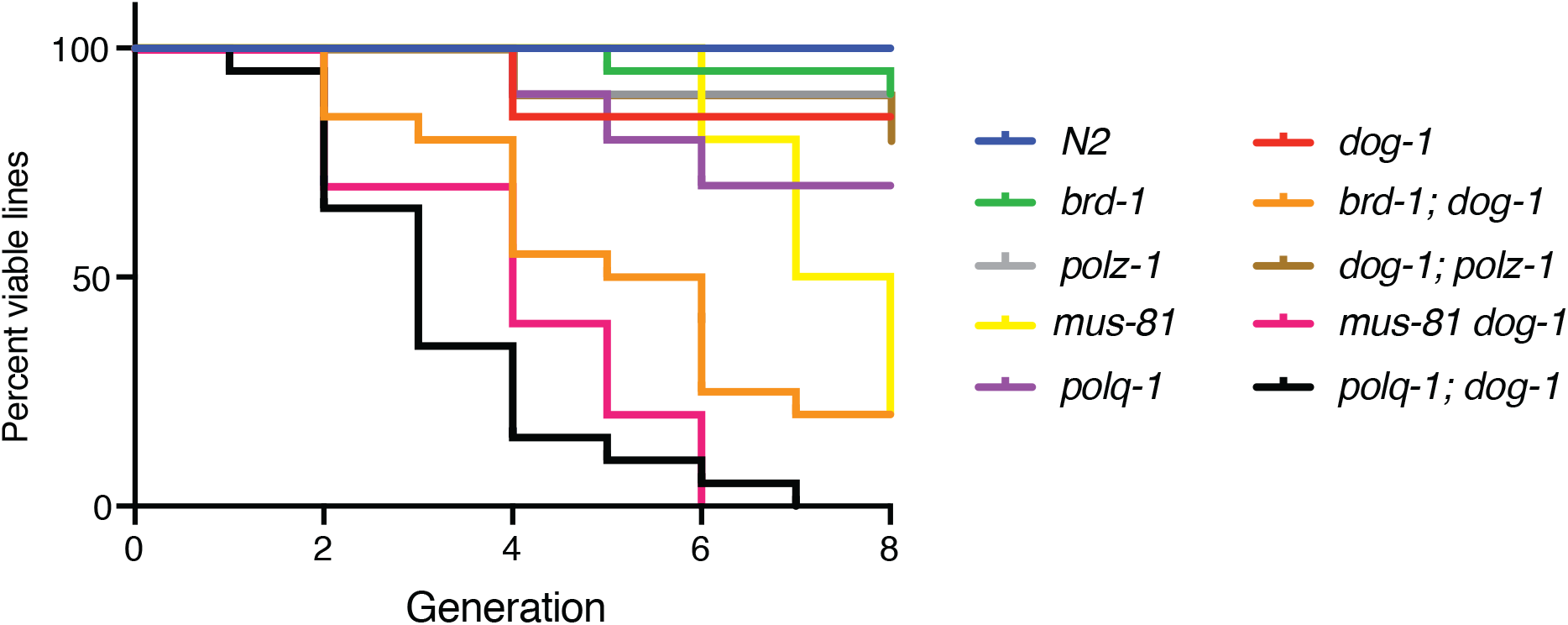
Effect of G-quadruplex stabilization on CX-5461-sensitive mutants. Multigenerational fitness assay. Loss of *polq-1, mus-81* or *brd-1* reduced the fitness of *dog-1* mutants.

## DISCUSSION

Key to the development of new anti-cancer therapeutic agents is understanding their off-target effects, mechanisms, and genotypic dependencies. While advances in target identification, chemical synthesis, and *in vitro* analysis have led to improvements in drug development, less progress has been made in improving toxicity and efficacy assays. The most common assay for mutagenicity is the bacteria-based Ames test [35], which has been used to assess the mutagenicity, photomutagenicity, and phototoxicity of chemotherapeutics [36]. The efficacy of the Ames test is limited because bacteria lack many of the genes responsible for the xenobiotic metabolism of drugs, have different DNA damage repair pathways than eukaryotes, and it only assays the reversion frequency of a single mutation. The small size, ease of handling, and powerful genetic tools of *C. elegans* provide a sophisticated *in vivo* toxicity assay that combines the technical advantages of a microorganism with greater biological complexity and a gene complement more akin to human. Furthermore, the cytochrome P450-based metabolic capabilities of *C. elegans* are broadly similar to those of mammals [37]. For these reasons, *C. elegans* has been used as an *in vivo* model system to predict the effect of chemicals on mammalian development [10], germline function [8,9], mutagenicity [13] and toxicity [38]. We have used a complementary suite of mutagenicity, mutational profile, and genotypic sensitivity assays that utilize *C. elegans* to characterize the new anticancer chemotherapeutic CX-5461.

Some anti-cancer drugs, such as vemurafenib, tamoxifen, and docetaxel, and many quinolonebased drugs can cause phototoxic reactions [22,39]. CX-5461, which contains a quinolone backbone, has resulted in photosensitivity in some patients [16,17]. We were able to phenocopy the photosensitivity in *C. elegans* and determine that the light sensitivity was accompanied by reactive oxygen species-mediated phototoxicity. We demonstrate that *C. elegans* can be used as an animal model to investigate drug-associated photosensitivity and test genetic and environmental factors affecting photosensitivity and resistance. Given the strong ROS-mediated phototoxicity and drug properties of CX-5461, CX-5461 may be useful for photodynamic anticancer therapy, in which targeted light is used to activate a photosensitizer within cancer cells leading to cell death.

*C. elegans* has proven to be an excellent model with which to investigate the mutagenicity and mutational profiles of DNA damage response mutants or genotoxic compounds [11–13]. CX5461 was mutagenic and the mutagenicity was increased by exposure to UVA light. The recessive mutation frequencies for CX-5461 and CX-5461 + UVA were comparable to exposure to 5 mM and 25 mM Ethyl Methane Sulfonate (EMS), a common alkylating mutagen, or 1000 and 2500 rads gamma radiation [15].

A mutational profile is a composite of the mutational events that occurred in a cell and can shed light on the mechanisms of action of DNA interacting compounds. Genetic balancers in *C. elegans* allow for the capture of a broad range of mutagenic events. CX-5461-treated genomes had a complex mutational profile with individual distinct mutational signatures that included both CNVs and SNVs. The nature of CX-5461-DNA lesions can be inferred from the mutational signature and the genes required for CX-5461 tolerance. For example, it is unlikely that CX-5461 generates interstrand crosslinks (ICLs) because loss of the key Fanconi Anemia pathway gene, *fcd-2,* did not result in CX-5461 sensitivity. Most CX-5461-exposed genomes contained both CNVs and SNVs. CNVs are indicative of DSB formation and repair. The major pathways for the repair of CX-5461-induced DSBs in *C. elegans* appear to be MMEJ and HDR. Simultaneous loss of both pathways resulted in hypersensitivity to CX-5461. Two of the most informative CX-5461-sensitive mutants are *rfs-1* and*polq-1.* RFS-1 mediates HDR at replication fork blocking lesions but not at IR-induced DSBs [40]. POLQ-1 promotes MMEJ mutagenic bypass of replication fork stalling lesions [12] and *dog-1-*induced G-quadruplexes [34]. This strongly suggests that CX-5461 does not cause DSBs directly but rather generates replication blocking lesions, which in turn can lead to breaks. This is further supported by the observation that *polq-1* and *rfs-1* and other genes required for the tolerance of CX-5461, such as *brd-1, smrc-1,* and *xpf-1* are also involved in the bypass or repair of replication blocking G-quadruplex structures that form in *dog-1* mutants [34,40–42] and are essential for the multigenerational survival of *dog-1* mutants (Figure 5). However, we observed very few G-quadruplex forming sequences in the regions flanking SNVs or CNV breakpoints and we demonstrated that CX-5461 can intercalate into non-G-quadruplex forming DNA sequences. Taken together, these data suggest that CX5461 results in DNA lesions or structures that can stall or collapse replication forks leading to DSBs even in the absence of G-quadruplexes.

CX-5461 and CX-5461 + UVA exposure resulted in a high frequency of SNVs. The CX-5461 AN mutation signature was similar to the mutational signature observed in human cancers that have been exposed to aristolochic acid [43,44]. However, the extended sequence context differed between CX-5461 (CATG) and aristolochic acid (T/CAG). Aristolochic acid results almost exclusively in A-T changes, whereas CX-5461 results in A-N changes. The A-T changes in aristolochic are dependent on the translesion polymerase polζ [45]. The high frequency of A-N SNVs, the presence of clustered multinucleotide mutations, and the CX-5461 hypersensitivity of translesion synthesis (TLS) mutants confirm that TLS is needed to bypass CX-5461-induced lesions.

How might CX-5461 trigger TLSr *In silico* analysis predicts that the pharmacophore of CX5461 can intercalate into the DNA sequence CGATCG [32] in a configuration similar to that of the antineoplastic agent ellipticine. When ellipticine intercalates into DNA, there is a slight unwinding of the ApT and a lengthening of the DNA [31], which could be consistent with the gel shifts we observed with DNA incubated with CX-5461. This distortion could make the ApT more prone to TLS-mediated mutagenesis either directly or through secondary reactions with the exposed adenine. Furthermore, both aristolochic acid and ellipticine can form covalent DNA adducts after reductive activation by cytochromes P450. It is possible that CX-5461 forms covalent adducts with DNA upon metabolic processing in *C. elegans.*

Overall, we found that CX-5461 shares many properties with ellipticine: both can intercalate into DNA [32], induce the formation reactive oxygen species [46], and inhibit RNA Pol I [18,32]. Ellipticine inhibits topoisomerase II and can form covalent DNA adducts [47]. These properties are consistent with the effects of CX-5461 on *C. elegans* but will require further experiments for confirmation. Ellipticine belongs to a family of promising anti-cancer therapeutics with a wide range of cellular effects similar to the anthracycline-based chemotherapeutics such as doxorubicin. However, ellipticines have failed in stage 1 or 2 clinical trials due to adverse side effects [32]. Based on the mechanistic similarities between ellipticine and CX-5461, it is possible that CX-5461 can elicit a similar response as ellipticine in tumour cells with fewer adverse side effects.

In summary, *C. elegans* is a powerful platform with which to interrogate the *in vivo* biological properties of both new and established anticancer therapeutic agents. The mutant panel we assembled and queried with DNA damaging agents provides valuable information about the types of damage generated by new DNA damaging therapeutics. From these data, we find that CX-5461 is a multimodal anticancer agent with mechanistic similarities to ellipticines and anthracyclines. This suggests that CX-5461 may be a more broadly applicable anticancer drug with a therapeutic range beyond HDR-deficient tumours.

## MATERIALS AND METHODS

### Strains and Culturing

Nematode strains were maintained as described previously [48]. The alleles used in this study were: *atm-1(tm5027), brd-1(dw1), rfs-1(ok1372), cku-80(ok861), lig-4(ok716), hsr-9(ok759), polq-1(tm2026), polh-1(lf31), polk-1(lf29), fcd-2 (tm1298), fan-1(tm423), fncm-1(tm3148), msh-2(ok2410), ercc-1(tm1981), xpa-1(ok698), mus-81(tm1937), rcq-5(tm424), rtel-1(tm1866), helq-1 (tm2134), dog-1(gk10), wrn-1(gk99), let-418(n3536), him-1(e879), hda-3(ok1991), gld-1(op236), cep-1(gk138), dvc-1(ok260), smrc-1(gk176502),* and *polz-1(gk919715).* Bristol N2 was used as wild type in all experiments. Some strains were provided by the CGC, which is funded by NIH Office of Research Infrastructure Programs (P40 OD010440), and some knockout alleles were provided by the Shohei Mitani laboratory. *smrc-1(gk176502), smrc-1(gk784642)*, and *polz-1(gk919715)* were Million Mutation Project alleles [29] provided by the Moerman lab and were outcrossed at least six times. Some strains were generated by the International *C. elegans* Gene Knockout Consortium [49] and by the National Bioresource Project of Japan.

### UVA irradiation

UVA Source: predominantly 365 nm, Black-Ray^®^ UV Bench Lamp (Model: XX-15L). Before each UVA exposure, the light source output was determined by a longwave ultraviolet measuring meter (Model: J-221). Different UVA exposures were achieved by varying the exposure times.

### Quantitative acute assay

Synchronized one-day-old adults were incubated in 100μM CX5461 (in NaH_2_PO_4_) diluted in M9 buffer containing OP50, carbenicillin (50 μg/ml) and 1X nystatin for ~18 hours. Following treatment, the animals were allowed to recover for 0.5 h on OP50 containing NGM plates before UVA irradiation (if applicable) and then plated at ten per plate in triplicates on NGM plates for a 4 h interval (18 to 22 h post-treatment), and then removed. The number of both arrested embryos and hatching larvae was counted one day later in order to calculate the percentage of embryo survival after treatment. All results were from at least 30 treated animals (3 plates with 10 animals per plate).

The sensitivity score was calculated by normalizing the embryo survival rate under drug-treated condition to non-drug condition with respect to that of wild-type N2 animals.

### Mutagenesis screen for CX-5461

Strain BC2200 *dpy-18/eT1(III);unc-46/eT1(V)* was used in the mutagenesis screen. *dpy-18/eT1(III);unc-46/eT1(V)* animals were treated with or without 100μM CX-5461 for 18 hours before UVA irradiation, and 200 single *dpy-18/eT1(III);unc-46/eT1(V)* F1s were picked in each condition. Sterile phenotype at F1 is considered as acquiring a dominant lethal mutation, and lines in which F2 or later generation that do not have Dpy Unc animals are counted as acquiring a recessive lethal mutation on balanced regions of chromosome III or V.

### Genome Sequencing

The lines that acquired recessive lethal mutations were maintained for at least three generations. Worms were rinsed off with deionized water and concentrated. Genomic DNA was purified using Puregene^®^ Core Kit A (Qiagen). DNA-seq was performed at the Novogene Bioinformatics Institute (Beijing, China). Sequence reads were mapped to the *C. elegans* reference genome version WS230 (http://www.wormbase.org) using the short-read aligner BWA [50], which resulted in an average sequencing depth for each sample ranging from 22x to 57x with a median of 34x. Single-nucleotide variants and small insertions/deletions were identified and filtered with the help of the SAMtools toolbox [51]. Candidate variants at genomic locations for which the parental <u>N2</u> strain had an agreement rate with the reference genome lower than 95% were eliminated from further consideration. Each variant was annotated with a custom Perl script and gene information downloaded from WormBase version WS230. Copy numbers were estimated from the alignments with a procedure analogous to that of Itani *et al.* [52] using 5 kb wide overlapping sliding windows with the alignments from the parental strain used as the reference.

### CX-5461 agarose gel shift and mung bean endonuclease assays

To test whether CX-5461 could intercalate into DNA, we incubated CX-5461 for one hour at room temperature with a PCR-generated dsDNA fragment and visualized the migration of DNA on a 1% agarose gel containing the dsDNA-specific dye SYBR-Safe. To determine whether CX-5461 affected mung bean endonuclease activity, we incubated a PCR product predicted to form a G quadruplex (G4) and a PCR product without a predicted G quadruplex (non-G4) with mung bean nuclease (MBN), which cleaves single-stranded or distorted double-stranded DNA, for one hour at the specified temperatures and assessed the endonuclease activity on a 1% Syber-Safe containing agarose gel.

### L1 exposure assay

Gravid animals were synchronized in the L1 stage by hypochlorite treatment (0.5M NaOH, 2% hypochlorite). After overnight starvation, approximately 100 L1 larvae of each mutant strain were incubated in 50 μl of M9 buffer containing OP50, carbenicillin (50 μg/ml), 1X nystatin, 500 μM NaH_2_PO_4_ with and without 100 μM CX5461 for ~18 hours. Following treatment, worms were allowed to recover for 0.5 h on OP50-containing NGM plates before they were irradiated with the indicated amount of UVA exposure. Animals were imaged after 4 days following UVA exposure with 20x using microscope.

### Reactive oxygen species (ROS) measurement with H_2_DCFDA

Adult worms treated in 100μM CX-5461 for 18 hours were added with 25 μM 2’, 7’ - dichlorodihydrofluorescein diacetate (H_2_DCFDA), and incubated for another hour in dark before initial fluorescence measurement by Microplate reader. After initial measurement, worms were irradiated with the indicated amount of UVA exposure, and then immediately sent for a second measurement [24].

### Generational survival assay

Animals were plated individually and maintained at room temperature. Starting with 20 separate lines at P_0_ generation, a single L4 stage animal was transferred to a fresh plate at each generation. A line was scored as unsustainable when the parent worm was either sterile or produced only dead embryos.

### Cell culture and treatment with CX5461

HCT116 and HT29 wild-type cells were obtained from American Type Culture Collection. Cells were cultured in McCoy’s 5A medium (Life Technologies) supplemented with 10% FBS at 37°C and 5% CO_2_. CX5461 was purchased from Selleck Chemicals. Cells were seeded in 96-well format (6 technical replicates) and after 24 hours, CX5461 (or DMSO) diluted in McCoy’s 5A medium was added to wells. 2 hours post incubation in the drug, cells were exposed to 50J/sqm UVA and allowed to grow for 4 to 5 days. Cells were fixed in 3.7% paraformaldehyde and stained with Hoechst 33342 before nuclei were counted on Cellomics Arrayscan VTI.

### Yeast assays

Wild-type (BY4742) and *rad52*Δ yeast strains were diluted from mid-log phase to OD_600_=0.01 in 200μl SC media +/− CX-5461 in 96-well plates. Cells were incubated for 3 hours +/− CX-5461 with constant shaking before UVA treatment and subsequent loading to a TECAN M200 plate reader. OD_600_ readings were measured every 30 minutes over a period of 24 hours and plates were shaken for 10 minutes before each reading. Strains were tested in 3 replicates per plate per condition and area under the curve (AUC) was calculated for each replicate. Strain fitness was defined as the AUC of each yeast strain relative to the AUC of the wild-type strain (without CX-5461 and UVA treatment) grown on the same plate.

## ACCESSION NUMBERS

The raw sequence data from this study have been submitted to the NCBI BioProject (http://www.ncbi.nlm.nih.gov/bioproject) under accession number PRJNA540967 and can be accessed from the Sequence Read Archive (SRA; https://www.ncbi.nlm.nih.gov/sra).

## SUPPORTING INFORMATION LEGENDS

Supplementary Figure 1. Coverage plot of CX-5461-induced genome rearrangements using 5 kb wide overlapping sliding windows.

Supplementary Figure 2. Coverage plot of CX-5461 + UVA-induced genome rearrangements using 5 kb wide overlapping sliding windows.

Supplemental Figure 3. Distribution of CX-5461-induced SNVs across all six chromosomes. Note the higher frequency of on the balanced chromosomes III and V.

Supplementary Table 1. Table of CX-5461-induced SNVs.

