## SUPPLEMENTAL FIGURES for "Characterization of the phototoxicity, chemigenetic profile, and mutational signatures of the chemotherapeutic CX-5461 in *Caenorhabditis elegans*"

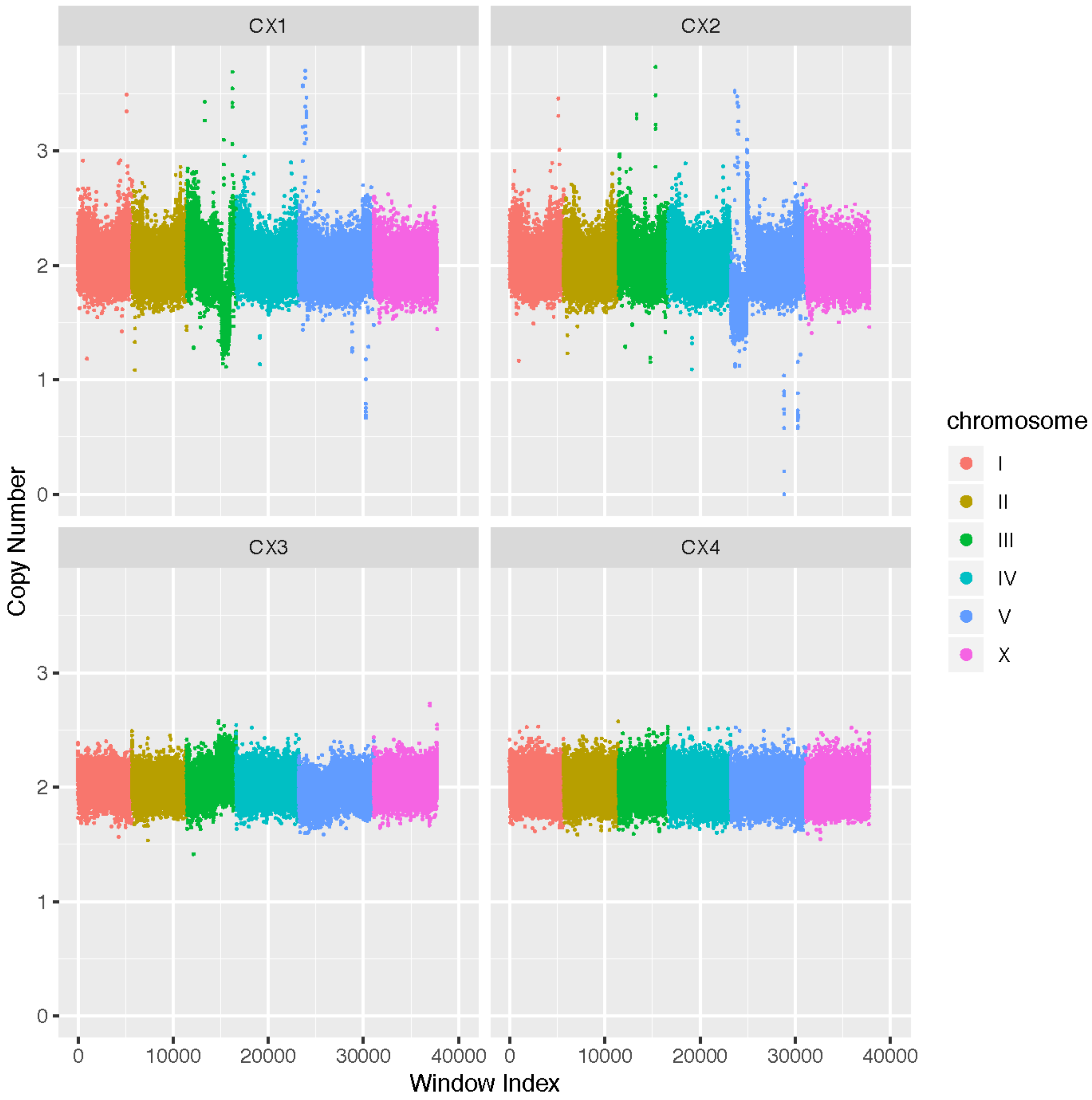


Supplementary Figure 1. Coverage plot of CX-5461-induced genome rearrangements using 5 kb wide overlapping sliding windows.

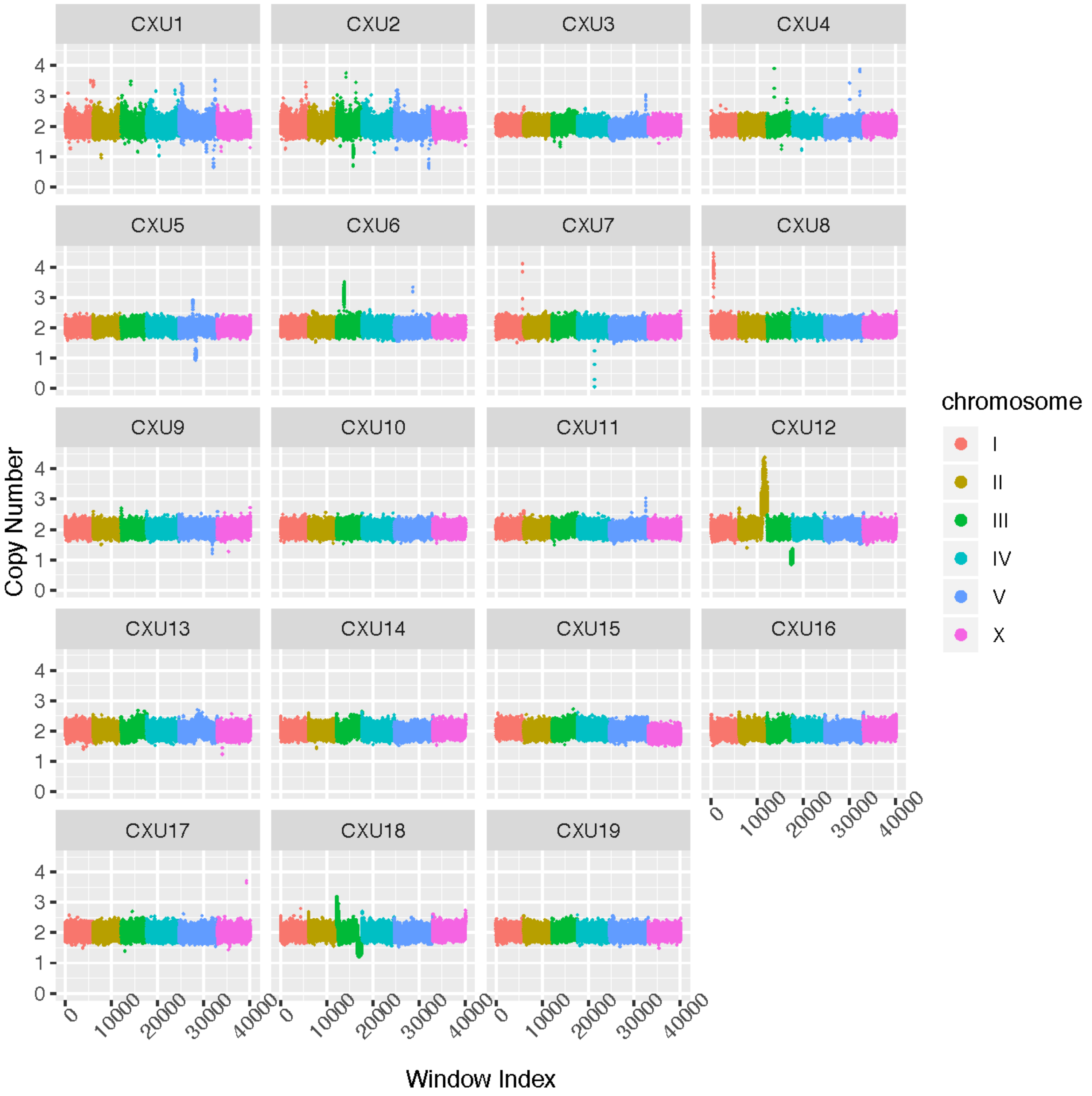


Supplementary Figure 2. Coverage plot of CX-5461 + UVA -induced genome rearrangements using 5 kb wide overlapping sliding windows.

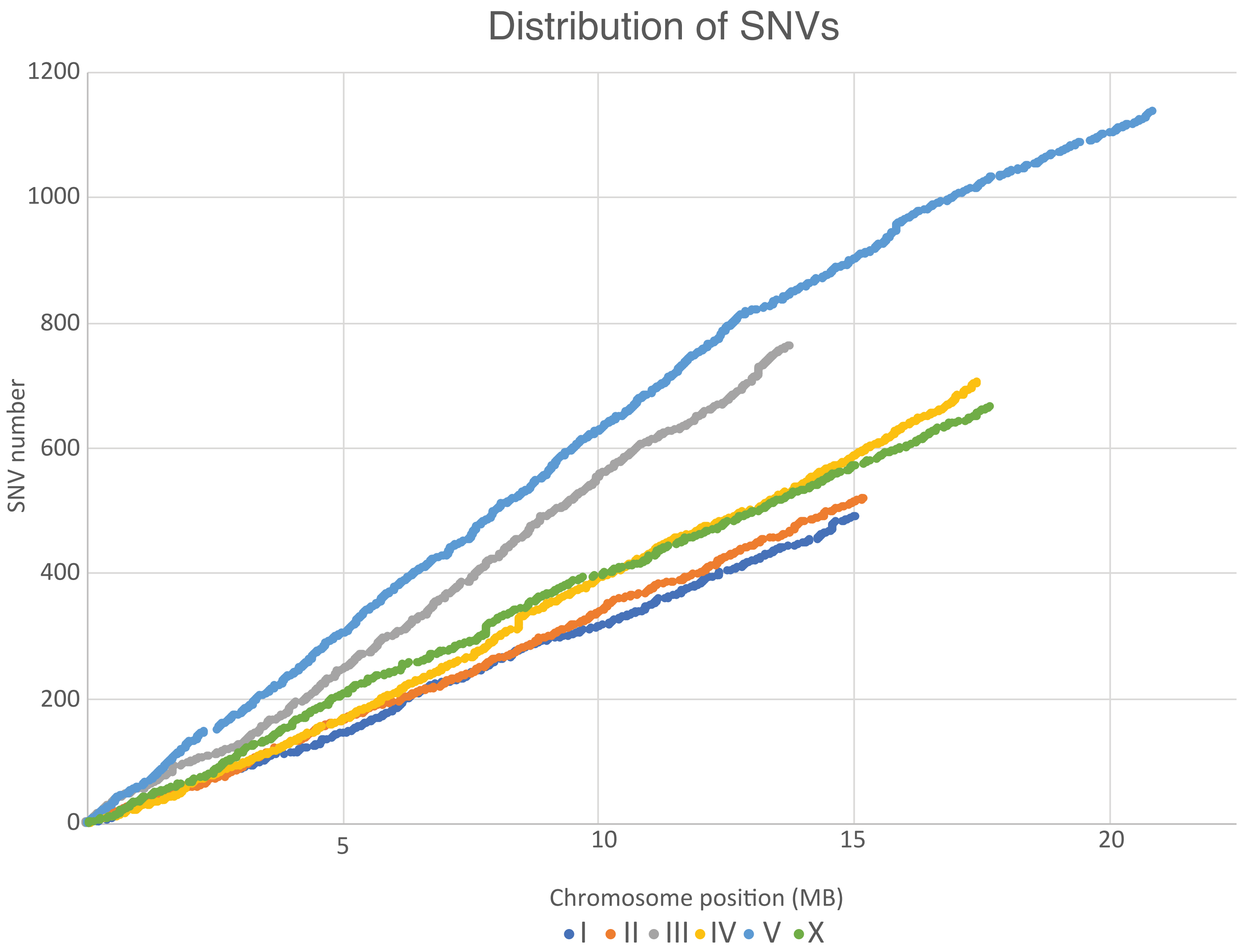


Supplemental Figure 3. Distribution of CX-5461-induced SNVs across all six chromosomes. Note the higher frequency of on the balanced chromosomes III and V.
